## Supplemental information for "Quantitative mechanistic model reveals key determinants of placental IgG transfer and informs prenatal immunization strategies"

### Appendix S1. Supporting information

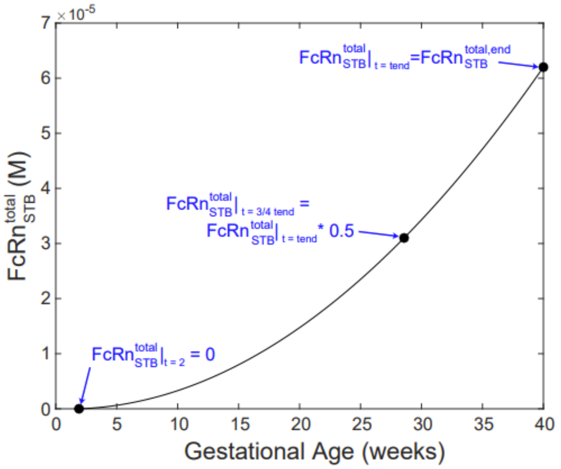


**Figure S1. Fc receptors are incorporated as dynamic offline variables to account for increasing expression across gestation.** $FcRn_{STB}$, $FcRn_{EC}$, and $Fc\gamma RIIb_{EC}$ were modeled as second order polynomials based on trends observed in rat placenta in a previous study by Wang et al. The fit for $FcRn_{STB}$ is shown here as an example. The trend of $FcRn_{STB}$ protein expression levels in rat placenta was scaled to the length of human gestation. The concentration of $FcRn_{STB}$ at full term was an optimized parameter. Additional constraints to achieve similar convex dynamics as observed by Wang et al are shown in blue. See equation 13 for more details.

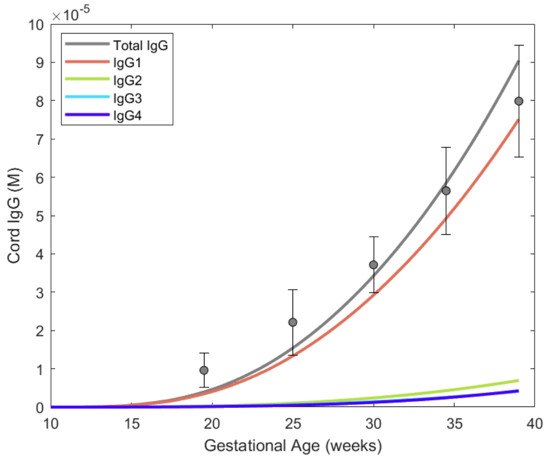


**Figure S2. The mechanistic model recapitulates bulk IgG transfer.** Bulk IgG in umbilical cord blood measured using cordocentesis in a cohort spanning gestational ages is shown as the cohort mean and standard deviation in grey circles. Simulated total fetal IgG and IgG subclasses across gestation are overlaid as solid curve. The curves for IgG3 and IgG4 are overlapping.

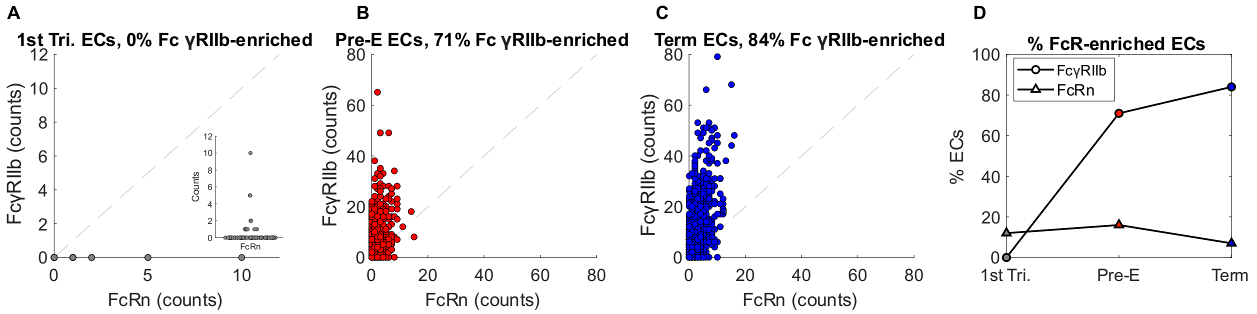


**Figure S3. Preferential expression of FCGR2B over FCGRT by placental endothelial cells increases across gestation. (A-C)** The scatter plots show the expression of FCGR2B and FCGRT in placental ECs in scRNA-seq data sets from three gestational time points: (A) first trimester, (B) early third trimester, and (C) late third trimester. Each dot corresponds to a single cell. Line y = x is indicated on each panel. In panel (A), a swarmchart of FcRn expression is included as an inlet to demonstrate the abundance of cells more clearly at the origin. **(D)** The trend of FcRn and FcgRIIb expression is shown across gestation as the percentage of ECs expressing FcgRIIb (circles) and FcRn (triangles) at each time point.

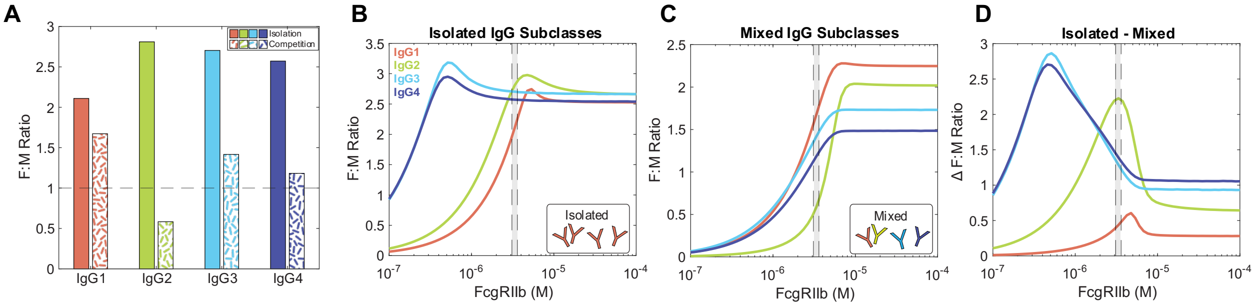


**Figure S4. IgG subclasses compete for FcgRIIb-mediated transcytosis in the placental mechanistic model. (A)** The bar plot represents the F:M ratio at parturition for a term pregnancy from independent simulations in the compartmental placental model with each subclass isolated (solid fill) or mixed with other IgG subclasses ("Competition", patterned fill). **(B-C)** The F:M ratio at parturition for a term pregnancy is shown for each subclass in isolation **(B)** or mixed with other subclasses **(C)** as in **(A)** over a range of FcgRIIb expression. **(D)** The difference (delta) in F:M ratio at parturition for a term pregnancy between isolation and competition conditions is shown over a range of FcgRIIb expression. In **(B-D),** the optimized range of FcgRIIb expression is indicated by the shaded region.

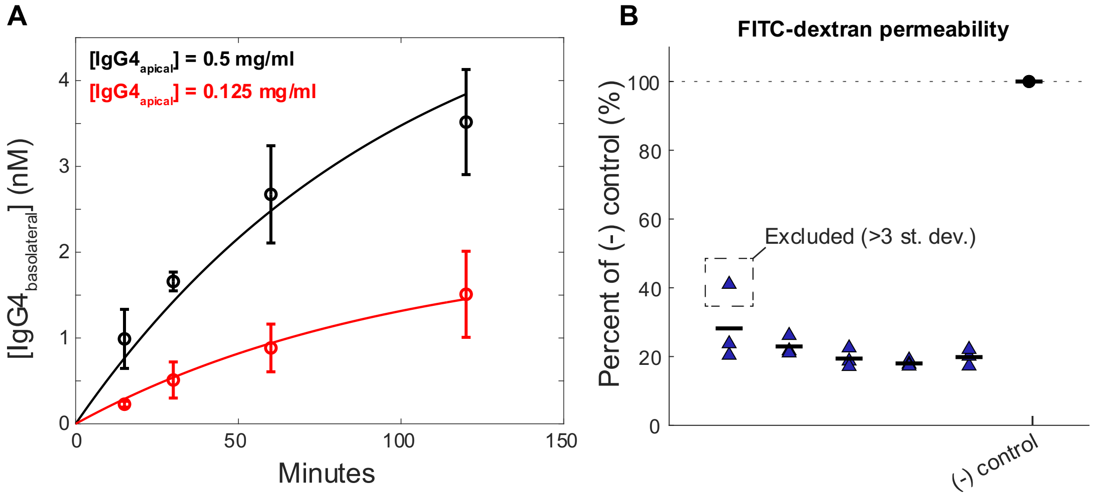


**Figure S5. Transwell simulation model parameters were fit to longitudinal *in vitro* data. (A)** Longitudinal dynamic data from IgG transcytosis measured *in vitro* in a HUVEC model are shown as the mean and standard deviation of 3 replicates. The overlaid curves show corresponding model simulations described in Methods. **(B)** The FITC-dextran permeability measurements demonstrates the criteria for rejecting a data point on the basis of faulty monolayer formation.

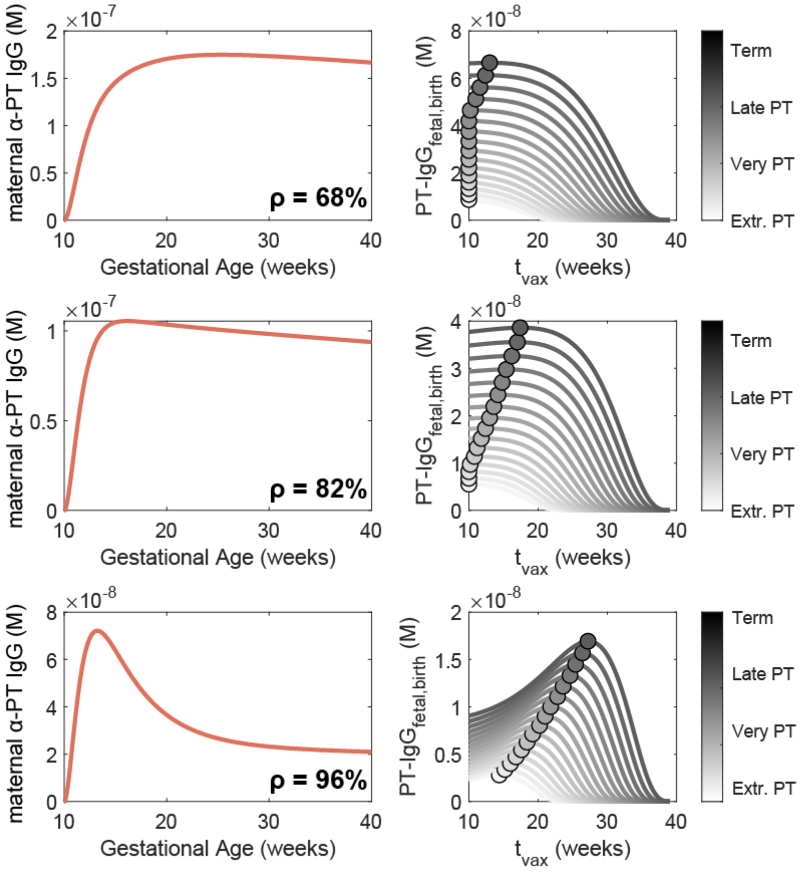


**Figure S6. The optimal immunization window during pregnancy depends on the existence of a transient spike in antigen-specific IgG.** The existence of a transient spike in anti-PT IgG is driven by parameter ρ, the proportion of short-lived antibody secreting cells. Three cases are shown within the range of optimized values of ρ reported in White et al. When ρ = 68% (top), the antibody response is driven by a larger proportion of long-lived antibody secreting cells resulting in a persistently high level of anti-PT IgG. Consequently, optimal immunization times were earlier in gestation for the majority of gestational age groups. When ρ = 82% (middle), the antibody response peaks and gradually declines and optimal immunization times were between 10-19 weeks gestation. When ρ = 96% (bottom), the spike in maternal IgG levels is highly transient and optimal immunization time scales with gestational age.

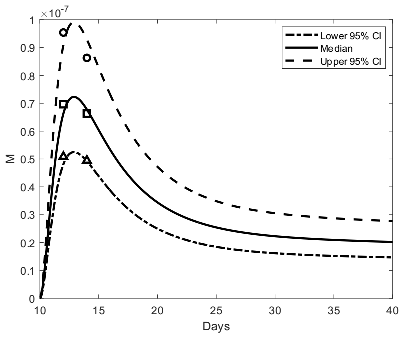


re

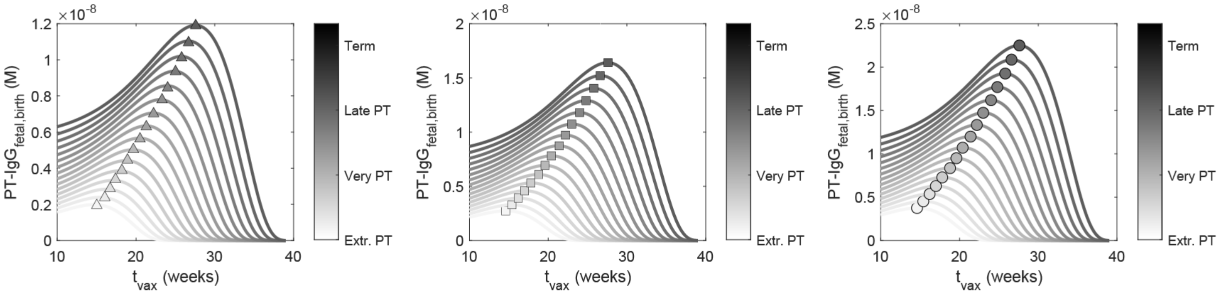


**Figure S7. The magnitude of the maternal anti-PT IgG response does not affect the predicted optimal immunization window.** Three models of the maternal anti-PT IgG response to Tdap immunization were fit to the median (squares) and upper (circles) and lower (triangles) 95% confidence intervals reported by van der Lee et al. The magnitude of the maternal anti-PT IgG spike affected the level of anti-PT IgG present in the fetus at the time of delivery but did not impact timing of the optimal immunization window.

### Placental IgG transfer model equations

Equations 1-4 represent IgG concentrations in the maternal plasma compartment:

$$\frac{dIgG1_M}{dt} = IgG1_M(k_{gen} - k_{up}) = 0 \quad (1)$$

$$\frac{dIgG2_M}{dt} = IgG2_M(k_{gen} - k_{up}) = 0 \quad (2)$$

$$\frac{dIgG3_M}{dt} = IgG3_M(k_{gen} - k_{up}) = 0 \quad (3)$$

$$\frac{dIgG4_M}{dt} = IgG4_M(k_{gen} - k_{up}) = 0 \quad (4)$$

Given the simplifying assumption that the overall concentration of IgG in the mother is constant such that the rate of new antibody generation ($k_{gen}$) and uptake by placental STBs ($k_{up}$) are equal. Equations 5-12 represent IgG in STB endosomes, either as monovalent IgG (5-8) or IgG in complex with FcRn in STB endosomes (9-12):

$$\frac{dIgG1_{STB}}{dt} = [k_{off}^{IgG1,FcRn} C_{STB}^{IgG1,FcRn} + k_{up} IgG1_M - k_{on}^{IgG1,FcRn} IgG1_{STB} FcRn_{STB}^{free} - k_{deg} IgG1_{STB}] \frac{1}{v_{STB}} \quad (5)$$

$$\frac{dIgG2_{STB}}{dt} = [k_{off}^{IgG2,FcRn} C_{STB}^{IgG2,FcRn} + k_{up} IgG2_M - k_{on}^{IgG2,FcRn} IgG2_{STB} FcRn_{STB}^{free} - k_{deg} IgG2_{STB}] \frac{1}{v_{STB}} \quad (6)$$

$$\frac{dIgG3_{STB}}{dt} = [k_{off}^{IgG3,FcRn} C_{STB}^{IgG3,FcRn} + k_{up} IgG3_M - k_{on}^{IgG3,FcRn} IgG3_{STB} FcRn_{STB}^{free} - k_{deg} IgG3_{STB}] \frac{1}{v_{STB}} \quad (7)$$

$$\frac{dIgG4_{STB}}{dt} = [k_{off}^{IgG4,FcRn} C_{STB}^{IgG4,FcRn} + k_{up} IgG4_M - k_{on}^{IgG4,FcRn} IgG4_{STB} FcRn_{STB}^{free} - k_{deg} IgG4_{STB}] \frac{1}{v_{STB}} \quad (8)$$

$$\frac{dC_{STB}^{IgG1,FcRn}}{dt} = [-k_{off}^{IgG1,FcRn} C_{STB}^{IgG1,FcRn} + k_{on}^{IgG1,FcRn} IgG1_{STB} FcRn_{STB}^{free} - k_{trans} C_{STB}^{IgG1,FcRn}] \frac{1}{v_{STB}} \quad (9)$$

$$\frac{dC_{STB}^{IgG2,FcRn}}{dt} = \left[-k_{off}^{IgG2,FcRn} C_{STB}^{IgG2,FcRn} + k_{on}^{IgG2,FcRn} IgG2_{STB} FcRn_{STB}^{free} - k_{trans} C_{STB}^{IgG2,FcRn}\right] \frac{1}{v_{STB}} \quad (10)$$

$$\frac{dC_{STB}^{IgG3,FcRn}}{dt} = \left[-k_{off}^{IgG3,FcRn} C_{STB}^{IgG3,FcRn} + k_{on}^{IgG3,FcRn} IgG3_{STB} FcRn_{STB}^{free} - k_{trans} C_{STB}^{IgG3,FcRn}\right] \frac{1}{v_{STB}} \quad (11)$$

$$\frac{dC_{STB}^{IgG4,FcRn}}{dt} = \left[-k_{off}^{IgG4,FcRn} C_{STB}^{IgG4,FcRn} + k_{on}^{IgG4,FcRn} IgG4_{STB} FcRn_{STB}^{free} - k_{trans} C_{STB}^{IgG4,FcRn}\right] \frac{1}{v_{STB}} \quad (12)$$

Equation 13 represents the concentration of free, unbound FcRn expressed within STB endosomes:

$$\begin{aligned}\frac{dFcRn_{STB}^{free}}{dt} = [&k_{off}^{IgG1,FcRn} C_{STB}^{IgG1,FcRn} + k_{off}^{IgG2,FcRn} C_{STB}^{IgG2,FcRn} + k_{off}^{IgG3,FcRn} C_{STB}^{IgG3,FcRn} \\ &+ k_{off}^{IgG4,FcRn} C_{STB}^{IgG4,FcRn} \\ &- \left(k_{on}^{IgG1,FcRn} IgG1_{STB} + k_{on}^{IgG2,FcRn} IgG2_{STB} + k_{on}^{IgG3,FcRn} IgG3_{STB} \right. \\ &\left. + k_{on}^{IgG4,FcRn} IgG4_{STB}\right) FcRn_{STB}^{free} \\ &+ k_{trans}\left(C_{STB}^{IgG1,FcRn} + C_{STB}^{IgG2,FcRn} + C_{STB}^{IgG3,FcRn} + C_{STB}^{IgG4,FcRn}\right) \\ &+ \Delta FcRn_{STB}^{total}] \frac{1}{v_{STB}} \quad (13)\end{aligned}$$

Where the total concentration of FcRn expressed by STBs increases exponentially across gestation, and the derivative of the second order polynomial curve fit ($\Delta FcRn_{STB}^{total}$) represents the increase in total FcRn at each time step:

$$FcRn_{STB}^{total} = at^2 + bt + c = FcRn_{STB}^{free} + [FcRn - \sum_i C_{STB}^{IgGi,FcRn}] \quad (13.i)$$

$$\Delta FcRn_{STB}^{total} = 2at + b \quad (13.ii)$$

Where parameters a, b, and c determined given the following conditions:

$$FcRn_{STB}^{total}|_{t=10} = 0$$

$$FcRn_{STB}^{total}|_{t=\frac{3}{4}t_{end}} = \frac{FcRn_{STB}^{total,end}}{2}$$

$$FcRn_{STB}^{total}|_{t=t_{end}} = FcRn_{STB}^{total,end}$$

And $FcRn_{STB}^{total,end}$ is a parameter optimized by CaliPro (Table 1). These assumptions are based on FcRn expression data in rat placenta scaled to the length of human gestation reported in Wang *et al* (31).

FcRn binding results in IgG transcytosis through STBs and into the intervillous stroma. IgG concentration in the stroma is represented by equations 14-17:

$$\frac{dIgG1_{STR}}{dt} = [k_{trans}C_{STB}^{IgG1,FcRn} + k_{off}^{IgG1,Fc\gamma RIIb}C_{EC}^{IgG1,Fc\gamma RIIb} - k_{on}^{IgG1,Fc\gamma RIIb}IgG1_{STR}Fc\gamma RIIb_{EC}^{free} - k_{up}IgG1_{STR}]\frac{1}{v_{STR}} \quad (14)$$

$$\frac{dIgG2_{STR}}{dt} = [k_{trans}C_{STB}^{IgG2,FcRn} + k_{off}^{IgG2,Fc\gamma RIIb}C_{EC}^{IgG2,Fc\gamma RIIb} - k_{on}^{IgG2,Fc\gamma RIIb}IgG2_{STR}Fc\gamma RIIb_{EC}^{free} - k_{up}IgG2_{STR}]\frac{1}{v_{STR}} \quad (15)$$

$$\frac{dIgG3_{STR}}{dt} = [k_{trans}C_{STB}^{IgG3,FcRn} + k_{off}^{IgG3,Fc\gamma RIIb}C_{EC}^{IgG3,Fc\gamma RIIb} - k_{on}^{IgG3,Fc\gamma RIIb}IgG3_{STR}Fc\gamma RIIb_{EC}^{free} - k_{up}IgG3_{STR}]\frac{1}{v_{STR}} \quad (16)$$

$$\frac{dIgG4_{STR}}{dt} = [k_{trans}C_{STB}^{IgG4,FcRn} + k_{off}^{IgG4,Fc\gamma RIIb}C_{EC}^{IgG4,Fc\gamma RIIb} - k_{on}^{IgG4,Fc\gamma RIIb}IgG4_{STR}Fc\gamma RIIb_{EC}^{free} - k_{up}IgG4_{STR}]\frac{1}{v_{STR}} \quad (17)$$

Stromal IgG binds to FcγRIIb on the EC surface. The concentration of IgG subclasses and FcγRIIb in complex formation on the EC surface is given by equations 18-21:

$$\frac{dC_{EC}^{IgG1,Fc\gamma RIIb}}{dt} = \left[-k_{off}^{IgG1,Fc\gamma RIIb}C_{EC}^{IgG1,Fc\gamma RIIb} + k_{on}^{IgG1,Fc\gamma RIIb}IgG1_{STR}Fc\gamma RIIb_{EC}^{free} - k_{trans}C_{EC}^{IgG1,Fc\gamma RIIb}\right]\frac{1}{v_{STR}} \quad (18)$$

$$\frac{dC_{EC}^{IgG2,Fc\gamma RIIb}}{dt} = \left[-k_{off}^{IgG2,Fc\gamma RIIb} C_{EC}^{IgG2,Fc\gamma RIIb} + k_{on}^{IgG2,Fc\gamma RIIb} IgG2_{STR} Fc\gamma RIIb_{EC}^{free} - k_{trans} C_{EC}^{IgG2,Fc\gamma RIIb}\right] \frac{1}{v_{STR}} \quad (19)$$

$$\frac{dC_{EC}^{IgG3,Fc\gamma RIIb}}{dt} = \left[-k_{off}^{IgG3,Fc\gamma RIIb} C_{EC}^{IgG3,Fc\gamma RIIb} + k_{on}^{IgG3,Fc\gamma RIIb} IgG3_{STR} Fc\gamma RIIb_{EC}^{free} - k_{trans} C_{EC}^{IgG3,Fc\gamma RIIb}\right] \frac{1}{v_{STR}} \quad (20)$$

$$\frac{dC_{EC}^{IgG4,Fc\gamma RIIb}}{dt} = \left[-k_{off}^{IgG4,Fc\gamma RIIb} C_{EC}^{IgG4,Fc\gamma RIIb} + k_{on}^{IgG4,Fc\gamma RIIb} IgG4_{STR} Fc\gamma RIIb_{EC}^{free} - k_{trans} C_{EC}^{IgG4,Fc\gamma RIIb}\right] \frac{1}{v_{STR}} \quad (21)$$

And the concentration of free FcγRIIb is given by equation 22:

$$\frac{dFc\gamma RIIb_{EC}^{free}}{dt} = [k_{off}^{IgG1,Fc\gamma RIIb} C_{EC}^{IgG1,Fc\gamma RIIb} + k_{off}^{IgG2,Fc\gamma RIIb} C_{EC}^{IgG2,Fc\gamma RIIb} + k_{off}^{IgG3,Fc\gamma RIIb} C_{EC}^{IgG3,Fc\gamma RIIb} + k_{off}^{IgG4,Fc\gamma RIIb} C_{EC}^{IgG4,Fc\gamma RIIb} - \left(k_{on}^{IgG1,Fc\gamma RIIb} IgG1_{STR} + k_{on}^{IgG2,Fc\gamma RIIb} IgG2_{STR} + k_{on}^{IgG3,Fc\gamma RIIb} IgG3_{STR} + k_{on}^{IgG4,Fc\gamma RIIb} IgG4_{STR}\right) Fc\gamma RIIb_{EC}^{free} + k_{trans}\left(C_{EC}^{IgG1,Fc\gamma RIIb} + C_{EC}^{IgG2,Fc\gamma RIIb} + C_{EC}^{IgG3,Fc\gamma RIIb} + C_{EC}^{IgG4,Fc\gamma RIIb}\right) + \Delta Fc\gamma RIIb_{EC}^{total}] \frac{1}{v_{STR}} \quad (22)$$

Where $\Delta Fc\gamma RIIb_{EC}^{total}$ is the derivative of the second order polynomial curve fit representing increasing FcγRIIb concentration across gestation:

$$Fc\gamma RIIb_{EC}^{total} = at^2 + bt + c \quad (22.i)$$

$$\Delta Fc\gamma RIIb_{EC}^{total} = 2at + b \quad (22.ii)$$

Where a, b, and c are parameters determined given a value FcγRIIb at term ($Fc\gamma RIIb_{EC}^{total,end}$) and similar constraints as described in equations (13.i – 13.ii). FcγRIIb binding results in IgG being taken up into EC endosomes where IgG dissociates from FcγRIIb and is trafficked across the ECs.

In contrast, EC FcRn binds IgG at an acidic pH within EC endosomes, which is modeled as:

$$\frac{dFcRn_{EC}^{free}}{dt} = [k_{off}^{IgG1,FcRn} C_{EC}^{IgG1,FcRn} + k_{off}^{IgG2,FcRn} C_{EC}^{IgG2,FcRn} + k_{off}^{IgG3,FcRn} C_{EC}^{IgG3,FcRn}$$

$$+ k_{off}^{IgG4,FcRn} C_{EC}^{IgG4,FcRn}$$

$$- \left(k_{on}^{IgG1,FcRn} IgG1_{EC} + k_{on}^{IgG2,FcRn} IgG2_{EC} + k_{on}^{IgG3,FcRn} IgG3_{EC}\right.$$

$$\left. + k_{on}^{IgG4,FcRn} IgG4_{EC}\right) FcRn_{EC}^{free}$$

$$+ k_{trans}\left(C_{EC}^{IgG1,FcRn} + C_{EC}^{IgG2,FcRn} + C_{EC}^{IgG3,FcRn} + C_{EC}^{IgG4,FcRn}\right)$$

$$+ \Delta FcRn_{EC}^{total}] \frac{1}{v_{EC}} \quad (23)$$

Where $\Delta FcRn_{EC}^{total}$ is the derivative of the second order polynomial curve fit representing increasing FcRn concentration across gestation (S1 Fig):

$$FcRn_{EC}^{total} = at^2 + bt + c \quad (23.i)$$

$$\Delta FcRn_{EC}^{total} = 2at + b \quad (23.ii)$$

Where a, b, and c are parameters determined given a value FcRn at term ($FcRn_{EC}^{total,end}$) and similar constraints as described in equations (13.i – 13.ii). The concentration of FcRn-IgG complexes within EC endosomes is given by equations 24-27:

$$\frac{dC_{EC}^{IgG1,FcRn}}{dt} = \left[-k_{off}^{IgG1,FcRn} C_{EC}^{IgG1,FcRn} + k_{on}^{IgG1,FcRn} IgG1_{EC} FcRn_{EC}^{free} - k_{trans} C_{EC}^{IgG1,FcRn}\right] \frac{1}{v_{EC}} \quad (24)$$

$$\frac{dC_{EC}^{IgG2,FcRn}}{dt} = \left[-k_{off}^{IgG2,FcRn} C_{EC}^{IgG2,FcRn} + k_{on}^{IgG2,FcRn} IgG2_{EC} FcRn_{EC}^{free} - k_{trans} C_{EC}^{IgG2,FcRn}\right] \frac{1}{v_{EC}} \quad (25)$$

$$\frac{dC_{EC}^{IgG3,FcRn}}{dt} = \left[-k_{off}^{IgG3,FcRn} C_{EC}^{IgG3,FcRn} + k_{on}^{IgG3,FcRn} IgG3_{EC} FcRn_{EC}^{free} - k_{trans} C_{EC}^{IgG3,FcRn}\right] \frac{1}{v_{EC}} \quad (26)$$

$$\frac{dC_{EC}^{IgG4,FcRn}}{dt} = \left[-k_{off}^{IgG4,FcRn} C_{EC}^{IgG4,FcRn} + k_{on}^{IgG4,FcRn} IgG4_{EC} FcRn_{EC}^{free} - k_{trans} C_{EC}^{IgG4,FcRn}\right] \frac{1}{v_{EC}} \quad (27)$$

The concentration of free IgG within EC endosomes is given by equations 28-32:

$$\frac{dIgG1_{EC}}{dt} = [k_{up}IgG1_{STR} - k_{on}^{IgG1,FcRn}IgG1_{EC}FcRn_{EC}^{free} + k_{off}^{IgG1,FcRn}C_{EC}^{IgG1,FcRn} - k_{deg}IgG1_{EC}]\frac{1}{v_{EC}} \quad (28)$$

$$\frac{dIgG2_{EC}}{dt} = [k_{up}IgG2_{STR} - k_{on}^{IgG2,FcRn}IgG2_{EC}FcRn_{EC}^{free} + k_{off}^{IgG2,FcRn}C_{EC}^{IgG2,FcRn} - k_{deg}IgG2_{EC}]\frac{1}{v_{EC}} \quad (29)$$

$$\frac{dIgG3_{EC}}{dt} = \left[k_{up}IgG3_{STR} - k_{on}^{IgG3,FcRn}IgG3_{EC}FcRn_{EC}^{free} + k_{off}^{IgG3,FcRn}C_{EC}^{IgG3,FcRn} - k_{deg}IgG3_{EC}\right]\frac{1}{v_{EC}} \quad (30)$$

$$\frac{dIgG4_{EC}}{dt} = \left[k_{up}IgG4_{STR} - k_{on}^{IgG4,FcRn}IgG4_{EC}FcRn_{EC}^{free} + k_{off}^{IgG4,FcRn}C_{EC}^{IgG4,FcRn} - k_{deg}IgG4_{EC}\right]\frac{1}{v_{EC}} \quad (31)$$

The IgG concentration in the fetus is represented by equations 32-35:

$$\frac{dIgG1_{F}}{dt} = [k_{trans}(C_{EC}^{IgG1,FcRn} + C_{EC}^{IgG1,Fc\gamma RIIb}) - \delta_{Ab}]\frac{1}{v_{F}} \quad (32)$$

$$\frac{dIgG2_{F}}{dt} = \left[k_{trans}(C_{EC}^{IgG2,FcRn} + C_{EC}^{IgG2,Fc\gamma RIIb}) - \delta_{Ab}\right]\frac{1}{v_{F}} \quad (33)$$

$$\frac{dIgG3_{F}}{dt} = \left[k_{trans}(C_{EC}^{IgG3,FcRn} + C_{EC}^{IgG3,Fc\gamma RIIb}) - \delta_{Ab}\right]\frac{1}{v_{F}} \quad (34)$$

$$\frac{dIgG4_{F}}{dt} = [k_{trans}(C_{EC}^{IgG4,FcRn} + C_{EC}^{IgG4,Fc\gamma RIIb}) - \delta_{Ab}]\frac{1}{v_{F}} \quad (35)$$

### Tdap immunization model equations

Equations 36-39 represent the naïve B cell response to maternal immunization against pertussis toxin. The concentration of antigen injected intramuscularly in the mother is given in equation 36:

$$\frac{dAg}{dt} = \begin{cases} t < t_{vax}, & 0 \\ t \geq t_{vax}, & [-\delta_{Ag}Ag]\frac{1}{v_M} \end{cases} \quad (36)$$

And is equal to 0 prior to $t_{vax}$. Equation 37 and 38 represent the concentration of short-lived antibody secreting cells ($S_{ASC}$) and long-lived antibody secreting cells ($L_{ASC}$) specific to pertussis toxin following immunization:

$$\frac{dS_{ASC}}{dt} = \begin{cases} t < t_{vax}, & 0 \\ t \geq t_{vax}, & [\rho k_{ASC} - \delta_{S-ASC}S_{ASC}]\frac{1}{v_M} \end{cases} \quad (37)$$

$$\frac{dL_{ASC}}{dt} = \begin{cases} t < t_{vax}, & 0 \\ t \geq t_{vax}, & [(1-\rho)k_{ASC} - \delta_{L-ASC}L_{ASC}]\frac{1}{v_M} \end{cases} \quad (38)$$

The concentration of IgG in the mother specific to pertussis toxin is represented by equation 39:

$$\frac{dIgG_M^{\alpha-PT}}{dt} = \begin{cases} t < t_{vax}, & 0 \\ t \geq t_{vax}, & \left[k_{IgG}(S_{ASC} + L_{ASC}) - \delta_{IgG}IgG_M^{\alpha-PT}\right]\frac{1}{v_M} \end{cases} \quad (39)$$

To perform pertussis toxin immunization simulations, equations 36-39 are layered into the model described by equations 1-35. New equations similar to 5-12, 14-21, and 24-35 are added to this set to represent anti-pertussis toxin IgG1, IgG2, IgG3, and IgG4 in STB endosomes, the stroma, EC endosomes, and the fetus. Rate-limiting equations 13, 22, and 23 are adjusted to account for competition between bulk IgG1-4 and anti-pertussis toxin IgG1-4 for $FcRn_{STB}$, $Fc\gamma RIIb_{EC}$, and $FcRn_{EC}$ binding for a total of 72 equations:

$$\frac{dFcRn_{STB}^{free}}{dt} = [k_{off}^{IgG1,FcRn}(C_{STB}^{IgG1,FcRn} + C_{STB,\alpha-PT}^{IgG1,FcRn}) + k_{off}^{IgG2,FcRn}(C_{STB}^{IgG2,FcRn} + C_{STB,\alpha-PT}^{IgG2,FcRn})$$

$$+ k_{off}^{IgG3,FcRn}(C_{STB}^{IgG3,FcRn} + +C_{STB,\alpha-PT}^{IgG3,FcRn}) + k_{off}^{IgG4,FcRn}(C_{STB}^{IgG4,FcRn} + +C_{STB,\alpha-PT}^{IgG4,FcRn})$$

$$- \left(k_{on}^{IgG1,FcRn}(IgG1_{STB} + IgG1_{STB}^{\alpha-PT}) + k_{on}^{IgG2,FcRn}(IgG2_{STB} + IgG2_{STB}^{\alpha-PT})\right.$$

$$\left. + k_{on}^{IgG3,FcRn}(IgG3_{STB} + IgG3_{STB}^{\alpha-PT}) + k_{on}^{IgG4,FcRn}(IgG4_{STB} + IgG4_{STB}^{\alpha-PT})\right)FcRn_{STB}^{free}$$

$$+ k_{trans}\left(C_{STB}^{IgG1,FcRn} + C_{STB}^{IgG2,FcRn} + C_{STB}^{IgG3,FcRn} + C_{STB}^{IgG4,FcRn} + C_{STB,\alpha-PT}^{IgG1,FcRn}\right.$$

$$\left. + C_{STB,\alpha-PT}^{IgG2,FcRn} + C_{STB,\alpha-PT}^{IgG3,FcRn} + C_{STB,\alpha-PT}^{IgG4,FcRn}\right) + \Delta FcRn_{STB}^{total}]\frac{1}{v_{STB}} \quad (40)$$

$$\frac{dFc\gamma RIIb_{STB}^{free}}{dt} = [k_{off}^{IgG1,Fc\gamma RIIb}(C_{EC}^{IgG1,Fc\gamma RIIb} + C_{EC,\alpha-PT}^{IgG1,Fc\gamma RIIb}) + k_{off}^{IgG2,Fc\gamma RIIb}(C_{EC}^{IgG2,Fc\gamma RIIb}$$

$$+ C_{EC,\alpha-PT}^{IgG2,Fc\gamma RIIb}) + k_{off}^{IgG3,Fc\gamma RIIb}(C_{EC}^{IgG3,Fc\gamma RIIb} + +C_{EC,\alpha-PT}^{IgG3,Fc\gamma RIIb})$$

$$+ k_{off}^{IgG4,Fc\gamma RIIb}(C_{EC}^{IgG4,Fc\gamma RIIb} + +C_{EC,\alpha-PT}^{IgG4,Fc\gamma RIIb})$$

$$- \left(k_{on}^{IgG1,Fc\gamma RIIb}(IgG1_{STR} + IgG1_{STR}^{\alpha-PT}) + k_{on}^{IgG2,Fc\gamma RIIb}(IgG2_{STR} + IgG2_{STR}^{\alpha-PT})\right.$$

$$+ k_{on}^{IgG3,Fc\gamma RIIb}(IgG3_{STR} + IgG3_{STR}^{\alpha-PT}) + k_{on}^{IgG4,Fc\gamma RIIb}(IgG4_{STR}$$

$$\left. + IgG4_{STR}^{\alpha-PT})\right)Fc\gamma RIIb_{EC}^{free}$$

$$+ k_{trans}\left(C_{EC}^{IgG1,Fc\gamma RIIb} + C_{EC}^{IgG2,Fc\gamma RIIb} + C_{EC}^{IgG3,Fc\gamma RIIb} + C_{EC}^{IgG4,Fc\gamma RIIb} + C_{EC,\alpha-PT}^{IgG1,Fc\gamma RIIb}\right.$$

$$\left. + C_{EC,\alpha-PT}^{IgG2,Fc\gamma RIIb} + C_{EC,\alpha-PT}^{IgG3,Fc\gamma RIIb} + C_{EC,\alpha-PT}^{IgG4,Fc\gamma RIIb}\right) + \Delta Fc\gamma RIIb_{EC}^{total}]\frac{1}{v_{EC}} \quad (41)$$

$$\frac{dFcRn_{EC}^{free}}{dt} = [k_{off}^{IgG1,FcRn}(C_{EC}^{IgG1,FcRn} + C_{EC,\alpha-PT}^{IgG1,FcRn}) + k_{off}^{IgG2,FcRn}(C_{EC}^{IgG2,FcRn} + C_{EC,\alpha-PT}^{IgG2,FcRn})$$

$$+ k_{off}^{IgG3,FcRn}(C_{EC}^{IgG3,FcRn} + +C_{EC,\alpha-PT}^{IgG3,FcRn}) + k_{off}^{IgG4,FcRn}(C_{EC}^{IgG4,FcRn} + +C_{EC,\alpha-PT}^{IgG4,FcRn})$$

$$-\left(k_{on}^{IgG1,FcRn}(IgG1_{EC} + IgG1_{EC}^{\alpha-PT}) + k_{on}^{IgG2,FcRn}(IgG2_{EC} + IgG2_{EC}^{\alpha-PT})\right.$$

$$\left. + k_{on}^{IgG3,FcRn}(IgG3_{EC} + IgG3_{EC}^{\alpha-PT}) + k_{on}^{IgG4,FcRn}(IgG4_{EC} + IgG4_{EC}^{\alpha-PT})\right)FcRn_{EC}^{free}$$

$$+ k_{trans}\left(C_{EC}^{IgG1,FcRn} + C_{EC}^{IgG2,FcRn} + C_{EC}^{IgG3,FcRn} + C_{EC}^{IgG4,FcRn} + C_{EC,\alpha-PT}^{IgG1,FcRn}\right.$$

$$\left. + C_{EC,\alpha-PT}^{IgG2,FcRn} + C_{EC,\alpha-PT}^{IgG3,FcRn} + C_{EC,\alpha-PT}^{IgG4,FcRn}\right) + \Delta FcRn_{EC}^{total}]\frac{1}{v_{EC}} \quad (42)$$
